# A Simple Method for Getting Standard Error on the Ratiometric Calcium Estimator

**DOI:** 10.1101/2020.12.22.424015

**Authors:** Simon Hess, Christophe Pouzat, Peter Kloppenburg

## Abstract

1.

The ratiometric fluorescent calcium indicator Fura-2 plays a fundamental role in the investigation of cellular calcium dynamics. Despite of its widespread use in the last 30 years, only one publication [2] proposed a way of obtaining confidence intervals on fitted calcium dynamic model parameters from single ’calcium transients’. Shortcomings of this approach are its requirement for a ’3 wavelengths’ protocol (excitation at 340 and 380 nm as usual plus at 360 nm, the isosbectic point) as well as the need for an autofluorence / background fluorescence model at each wavelength. We propose here a simpler method that eliminates both shortcommings:

a precise estimation of the standard errors of the raw data is obtained first,
the standard error of the ratiometric calcium estimator (a function of the raw data values) is derived using both the propagation of uncertainty and a Monte-Carlo method.

Once meaningful standard errors for calcium estimates are available, standard errors on fitted model parameters follow directly from the use of nonlinear least-squares optimization algorithms.

**Graphical abstract:** Figure 1:
How to get error bars on the ratiometric calcium estimator? The figure is to be read clockwise from the bottom right corner. The two measurements areas (region of interest, ROI, on the cell body and background measurement region, BMR, outside of the cell) are displayed on the frame corresponding to one actual experiment. Two measurements, one following an excitation at 340 nm and the other following an excitation at 380 nm are performed (at each ’time point’) from each region. The result is a set of four measures: adu340 (from the ROI), adu340B (from the BMR), adu380 and adu380B. *These measurements are modeled as realizations of Gaussian random variables*. The fact that the measurements as well as the subsequent quantities derived from them are random variable realization is conveyed throughout the figure by the use of Gaussian probability densities. The densities from the MRB are ’tighter’ because there are much more pixels in the MRB than in the ROI (the standard deviations of the densities shown on this figure have been enlarged for clarity, but their relative size has been preserved, the horizontal axis in black always starts at 0). The key result of the paper is that the standard deviation of the four Gaussian densities corresponding to the raw data (bottom of the figure) can be reliably estimated from the data alone, 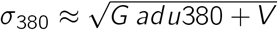, where *V* is the product of the CCD chip gain squared by the number of pixels in the ROI by the CCD chip readout variance. The algebric operations leading to the estimator (top right) are explicitely displayed. The paper explains how to compute the standard deviation of the derived distributions obtained at each step of the calcium concentration estimation.

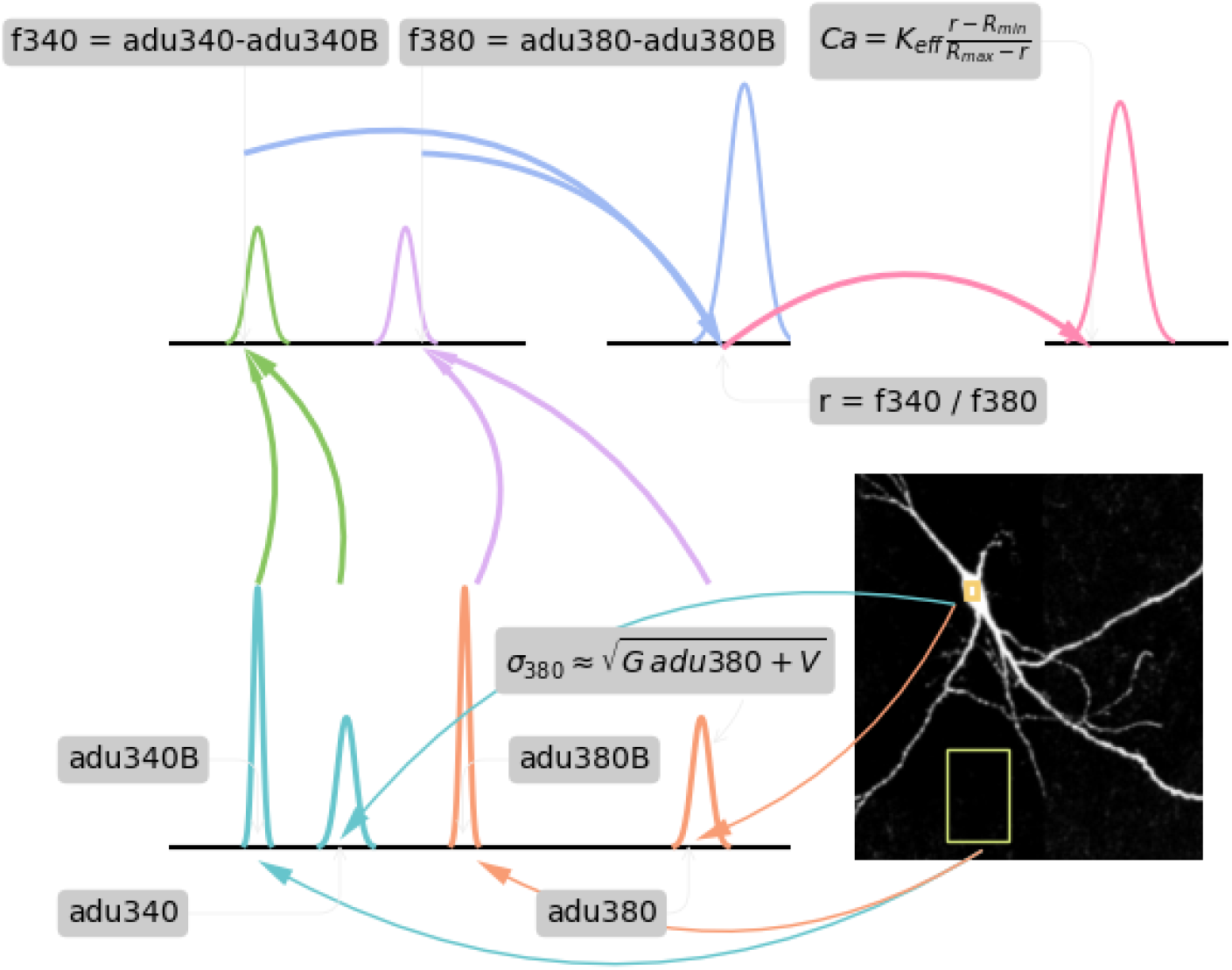

**Method name:** Standard error for the ratiometric calcium estimator

## 3. Method overview

### 3.1. Rational

Since its introduction by [1], the ratiometric indicator Fura-2 has led to a revolution in our understanding of the role of calcium ions (Ca^2+^) in neuronal and cellular function. This indicator provides a straightforward estimation of the free Ca^2+^ concentration ([Ca^2+^]) in neurons and cells with a fine spatial and time resolution. The experimentalist must determin a ’region of interest’ (`ROI`) within which the [Ca^2+^] can be assumed uniform and is scientifically relevant. Fluorescence must be measured following excitation at two different wavelengths: typically around 340 and 380 nm; and, since cells exhibit autofluorescence or ’background fluorescence’ at those wavelengths, the measured fluorescence intensity is made of two sources: the Fura-2 linked fluorescence and the autofluorescence. The measured intensity within the ROI is therefore usually corrected by subtrating from it an estimation of the autofluorescence intensity obtained from simultaneous measurements from a ’background measurement region’ (`BMR`); that is, a nearby region where there is no Fura-2. At a given time the experimentalist will therefore collect a fluorescence intensity measurement from the ROI at 340 and 380 nm; we are going to write `adu340` and `adu380` these measurements, where ’adu’ stands for ’analog to digital unit’ and correspond to the raw output of the fluorescence measurement device, most frequently a charge-coupled device (`CCD`). The experimentalist will also collect intensity measurements from the BMR, measurements that we are going to write `adu340B` and `adu380B`. If `P CCD` pixels make the ROI and `P_B` pixels make the BMR and if the illumination time at 340 nm is T_340, while the illumination time at 380 nm is `T_380` (both times are measured in *s*), the experimentalist starts by estimating the fluorescence intensity per pixel per time unit following an excitation by a light flash of wavelengths *λ* (*λ* = 340 or 380 nm) as:

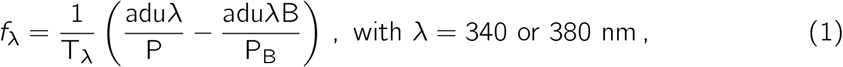

where an assumption of autofluorescence uniformity is implicitly made. The following ratio is then computed:

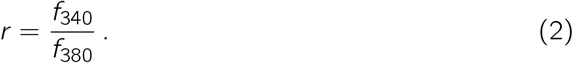

This is an important and attractive feature of the method as well as the origin of its name. Since only ratios are subsequently used, geometric factors like the volume of the Fura loaded region under the ROI do not need to be estimated.

The *estimated* [Ca^2+^] that we will write 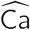 for short (the 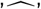 sign is used for marking estimated values) is then obtained, following [1, Eq. 5, p. 3447], with:

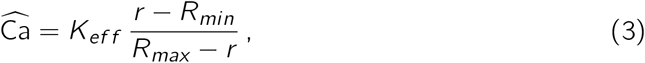

where *K_ef f_* (measured in *µM*), *R_min_* and *R_max_* are calibrated parameters (the last two parameters are ratios and are dimensionless).

If we now want to rigorously fit [Ca^2+^] dynamics models to sequences of 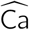, we need to get *standard errors*, 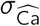, on our estimates. This is where the ratiometric method gets ’more involved’, at least if we want standard errors from a single transient as opposed to a mean of many transients. Despite the ubiquity of ratiometric measurements in neuroscience and cell physiology, we are aware of a single paper–by some of us [2]–where the ’standard error question’ was directly addressed. The method proposed in [2] requires a 3 wavelengths protocol: measurements at 340, 380 *and* 360 (the isosbestic wavelength) nm; it drops, so to speak, the above advantage of working with a ratiometric estimator since it fits directly the adu340 and adu380 data (at the cost of estimating some geometry related parameters) and it requires a model of the autofluorescence dynamics if the latter is not stationnary. It therefore requires a slightly more complicated ’3 wavelengths’ recording protocol as well as a more involved fitting procedure. The dataset of the companion paper exhibits a clear but reversible autofluorescence rundown that cannot be ignored since autofluorescence accounts for half of the signal in the ROI. Rather that constructing / tailoring the accurate enough autofluorence models required by the ’direct approach’ of [2] we looked for an alternative method providing standard errors for the ratiometric estimator.

## 4. Ratiometric estimator variance

### 4.1. Fluorescence intensity

As detailed in [1, 2], the fluorescence intensities giving rise to the adu340, adu340B, adu380 and adu380B signals can be written as:

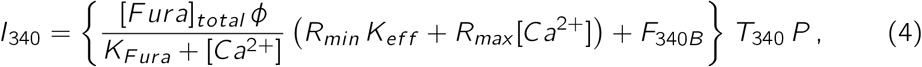

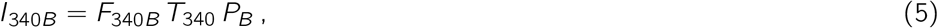

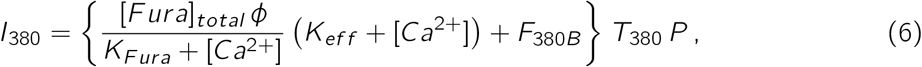

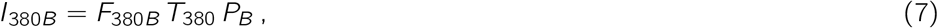

where *F_λB_* is the autofluorescence intensity per pixel per time unit at wavelength *λ*, *K_F ura_* is the Fura dissociation constant (a calibrated parameter measured in *µM*), [*F ura*]_*total*_, is the total (bound plus free) concentration of Fura in the cell (measured in *µM*) and *ϕ* is an experiment specific parameter (measured in 1*/µM/s*) lumping together the quantum efficiency, the neurite volume, etc (see [2] for details). When a ’3 wavelengths protocol’ is used (as was the case in the companion paper), it is easy to implement a self-consistency check based on these equations as explained in Sec. A.

### 4.2. Recorded signals adu340, adu340B, adu380 and adu380B

As detailed and discussed in [7, 2], the signal adu*λ* recorded with a CCD chip whose gain is *G* and whose read-out variance is 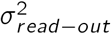 can be modeled as the realization of a Gaussian random variable ADU*λ* with parameters:

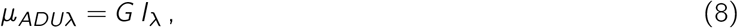

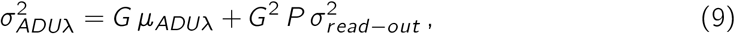

with the obvious adaptation when dealing with the BMR signal: *I_λ_* is replaced by *I_λB_* and *P* is replaced by *P_B_*.

### 4.3. Variance estimates for adu340, adu340B, adu380 and adu380B

So, to have the variance of *ADU_λ_* we need to know *I_λ_* and for that we need to know [*Ca*^2+^] (Eq. 4 and 6) precisely what we want to estimate. But the expected value of *ADU_λ_* is *G I_λ_* (Eq. 8), we can therefore use as a first approximation the observed value *adu_λ_* of *ADU_λ_* as a guess for *G I_λ_*, so in Eq. 9 we plug-in *adu_λ_* for *G I_λ_*, leading to:

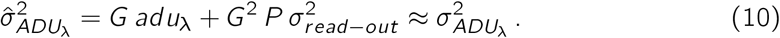

In other words, we will use the observed *adu_λ_* as if it were the actual fluorescence intensity times the CCD chip gain, *ADU_λ_* = *G I_λ_*, in order to estimate the variance. In doing so we will sometime slightly underestimate the actual variance (when the observed *adu_λ_* turns out to be smaller than *ADU_λ_*) and sometime slightly overestimate it (when the observed *adu_λ_* turns out to be larger than *ADU_λ_*). Since we are going to combine many such approximations, we expect–and we will substantiate this claim in Sec. 5–that overall the under-estimations will be compensated for by the over-estimations.

### 4.4. Variance estimate for 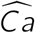

Now that we have a 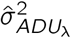 we can work with-that is, an estimate from the data alone—, we want to get 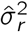 (Eq. 2) and 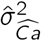. We can either use the propagation of uncertainty (also referred to as *error propagation*, *compounding of errors* or *delta method*) [6, 8] together with Eq. 2 and 3, or a ’quick’ Monte Carlo approach. We drop any explicit time index in the sequel in order to keep the equations more readable, but it should be clear that such variance estimates have to be obtained for each sampled point.

#### 4.4.1. Propagation of uncertainty

This method requires, in the general case, an assumption of ’small enough’ standard error since it is based on a first order Taylor expansion (see Sec. B for details). It leads first to the following expression for the variance, 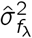, of *f_λ_* in Eq. 1:

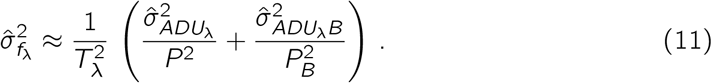

The variance 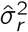 of *r* in Eq. 2 is then:

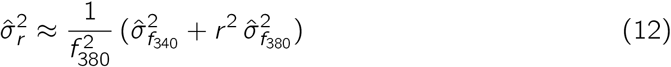

and the variance 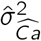 of 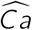 in Eq. 3 is:

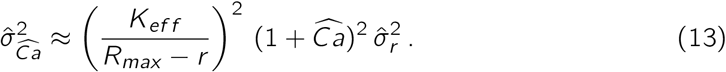

#### 4.4.2. Monte-Carlo method

Here we draw, *k* quadruple of vectors

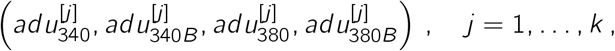

from four independent Gaussian distributions of the general form:

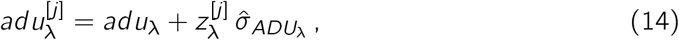

where *adu_λ_* is the observed value and 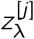 is drawn from a standard normal distribution. We then plug-in these quadruples into Eq. 1 leading to *k* couples:

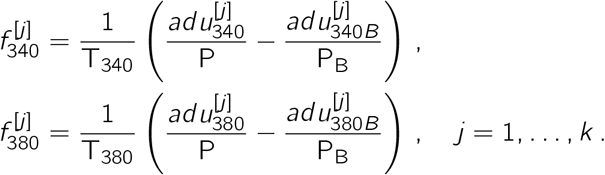

These *k* couples are ’plugged-in Eq. 2’ leading to *k r* ^[*j*]^:

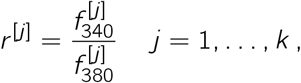

before plugging in the latter into Eq. 3 to get 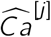:

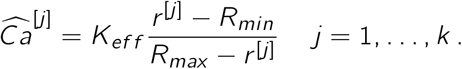

The empirical variance of these simulated observations will be our 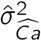:

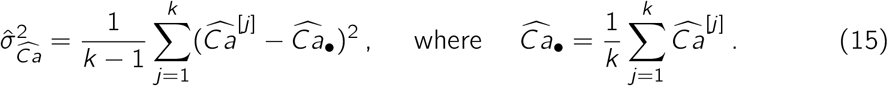

Since the Monte-Carlo method requires milder assumptions (the variances do not have to be small) and is easy to adapt, we tend to favor it, but both methods are working.

### 4.5. Comment

The present approach based on a 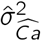 estimation is slightly less rigorous than the ’direct approach’ of [2] but it is far more flexible since it does not require an independent estimation / measurement of [*Fura*]_*total*_. In the companion paper we also chose to consider the calibrated parameters *K_ef f_*, *R_min_* and *R_max_* as known, while only an estimation (necessarily imprecise since the calibration procedure is subject to errors) is known. But the same batch of Fura was used for all experiments that should then all exhibit the same (systematic) error. Since in the companion paper, we are not trying to get the exact standard error of the calcium dynamics parameters but to show difference between two protocols (’classical whole-cell’ versus beta-escin perforated whole-cell), ignoring the uncertainty on the calibrated parameters makes our estimates less variable and helps making the difference, if it exists, clearer. For a true calcium buffering capacity study, the errors on the calibrated parameters should be accounted for and it would be straightforward to do it with our approach, simply by working out the necessary derivatives if one works with the propagation of uncertainty method, or by drawing also *K_ef f_*, *R_min_* and *R_max_* values from Gaussian distributions centered on their respective estimates with a SD given by the standard errors if one works with the Monte-Carlo approach.

## 5. Empirical validation

### 5.1. Rational

Equations 4, 5, 6, 7, together with Eq. 8 and 9 can be viewed as a data generation model. This means that if we choose model parameters values as well as an arbitrary [Ca^2+^] time course, we can simulate measurements (adu) at both wavelengths in the ROI as well as in the BMR. We can then use these simulated adu exactly as we used the actual data, namely get *r* (*t_i_*) (Eq. 2) and 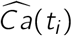 (Eq. 3) as well as the (squared) standard errors 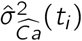 (Sec. 4.4).

Now if the 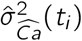 are good approximations for the actual but unknown 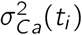, the distribution of the normalized residuals:

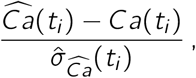

should be very close to a standard normal distribution. *This is precisely what we are going to check*.

### 5.2. Simulated data

We are going to use the first transient of dataset `DA_121219_E1` of the companion paper. The ’static’ parameters–that is the parameters not link to the calcium dynamics–used for the simulation are the actual experimental parameters rounded to the third decimal (Table 1).

**Table 1:** ’Static’ parameters used for the simulation.

| Parameter | Value |
| --- | --- |
| $R_{min}$ | 0.147 |
| $R_{max}$ | 1.599 |
| $K_{eff}$ | 1.093 ($\mu M$) |
| $K_{Fura}$ | 0.225 ($\mu M$) |
| $[Fura]_{total}\phi$ | 1.89e+05 ($s^{-1}$) |
| $T_{340}$ | 0.01 (s) |
| $T_{380}$ | 0.003 (s) |
| P | 3 |
| $P_B$ | 448 |
| G | 0.146 |
| $\sigma_{read-out}^2$ | 268.96 |
| $F_{340B}$ | 189512 ($s^{-1}$) |
| $F_{380B}$ | 711589 ($s^{-1}$) |

The simulated calcium dynamics is a monoexponential decay mimicking the tail of the transient:

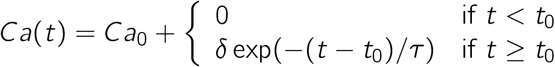

and the parameter values (Table 2) are just a rounded version of the fitting procedure output (see companion paper).

**Table 2:**
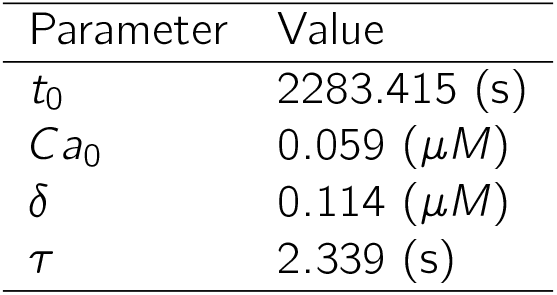
Calcium dynamics parameters used for the simulation. Time 0 is when seal is obtained.

| Parameter | Value |
| --- | --- |
| $t_0$ | 2283.415 (s) |
| $Ca_0$ | 0.059 ($\mu M$) |
| $\delta$ | 0.114 ($\mu M$) |
| $\tau$ | 2.339 (s) |

The simulated data obtained in that way are shown on Fig. 2 (blue traces) together with the actual data (red curves) they are supposed to mimic. At a qualitative level at least, our data generation model is able to produce realistic looking simulations.

**Figure 2:**
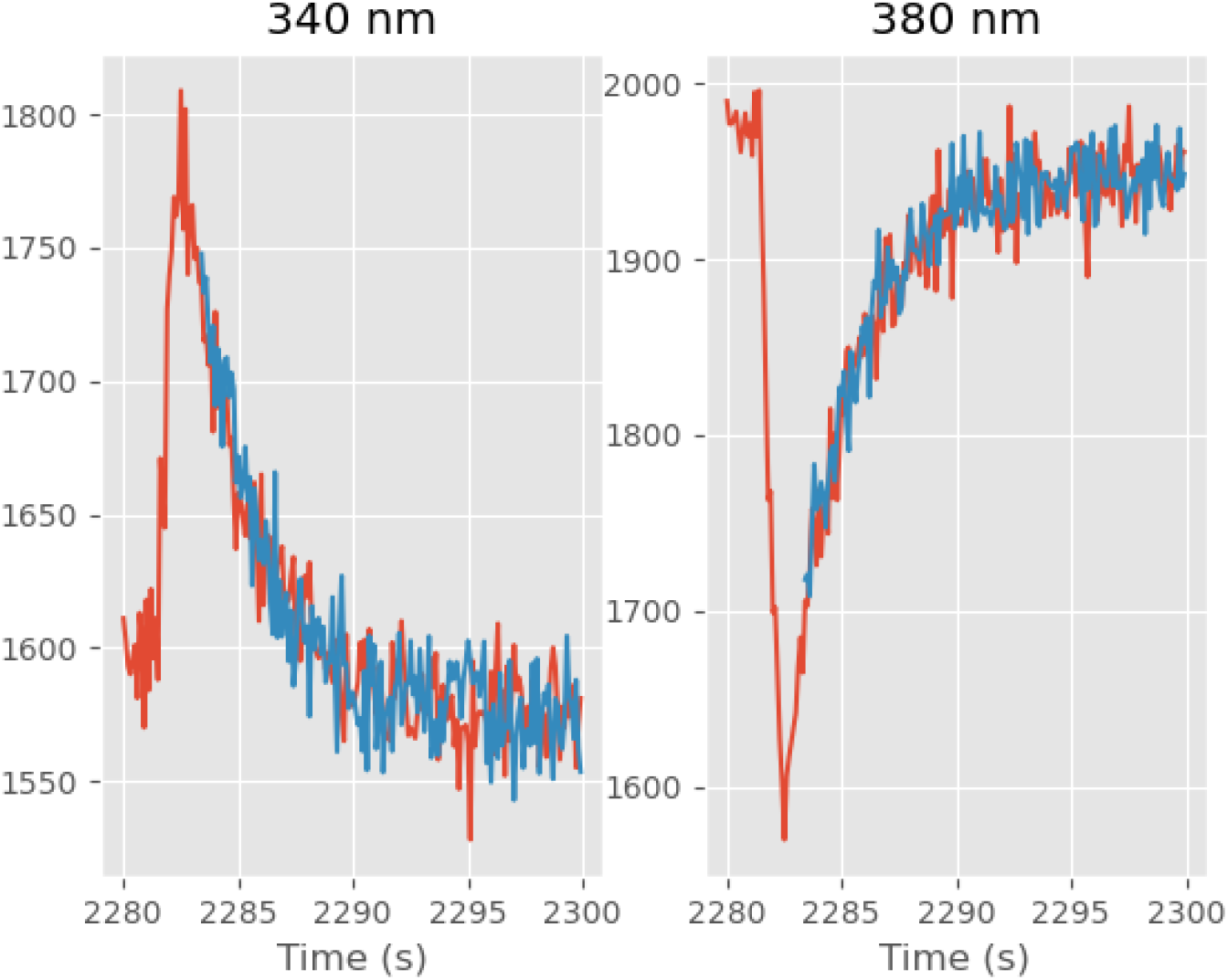
Observed (red) and simulated (blue) ADU at 340 (left) and 380 nm (right) for the first transient (only the late phase of the transient was simulated).

### 5.3. Software and simulation details

The methodological details of the measurements to which the analysis presented in the present manuscript was applied are described in the companion paper.

The simulations, computations and figures of the present manuscript were done with Python 3 (https://www.python.org/), numpy (https://numpy.org/), scipy and matplotlib (https://matplotlib.org/). The Python codes and the data required to reproduce the simulations and figures presented in this manuscript can be downloaded from GitLab (https://gitlab.com/c_pouzat/getting-se-on-ratiometric-ca-estimator).

The use of scipy was kept to a bar minimum to maximize code lifeduration (scipy tends to evolve too fast with minimal concern for backward compatibility). The random number generators used were therefore the ones of numpy: the uniform random number generator derives from the Permuted Congruential Generator (64-bit, PCG64) (https://www.pcg-random.org/) [5] while the normal random number generator is an adaptation of the Ziggurat method [3] of Julia (https://docs.julialang.org/en/v1/); unfortunately one has to check the source code of both numpy and Julia to find that out.

### 5.4. Are the standard errors of ratiometric estimator accurate?

Since the two 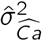 estimation methods, propagation of uncertainty and Monte-Carlo, agree at each time point within 2%, we illustrate in this section the results obtained with the Monte-Carlo method.

We take next the simulated data (blue curves on Fig. 2) together with the simulated background signals (not shown) as if they were actual data and we compute the ratiometric estimator and its standard error as described in Sec. 4.4, using *k* = 10^4^ replicates. Figure 3 shows the standardized residuals as well as the simulated data together with the true [Ca^2+^], we know it since we used it to simulate the data!

**Figure 3:**
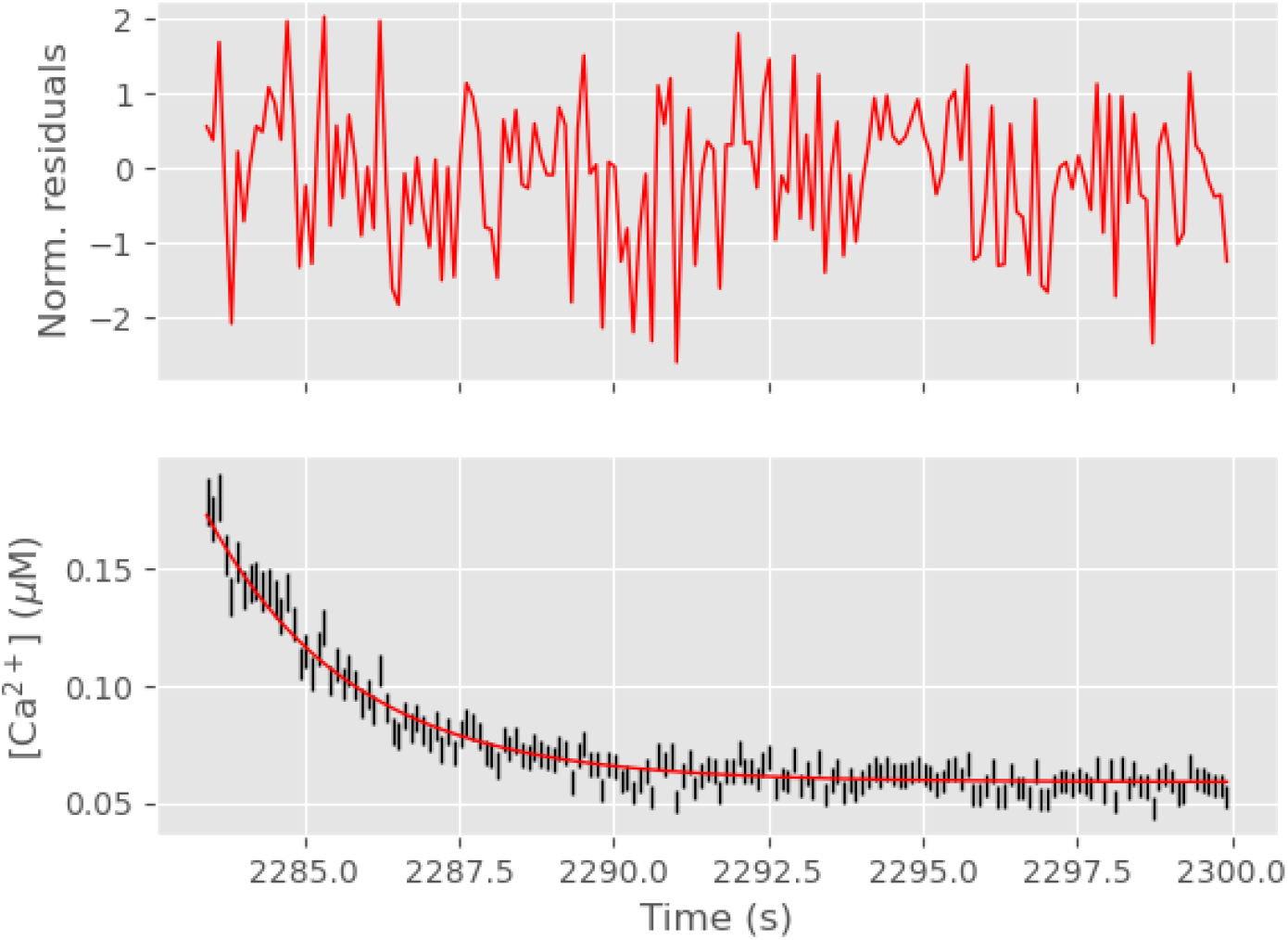
Top: Simulated ratiometric estimator - ’actual’ [Ca^2+^] divided by ratiometric estimator standard error (if everything goes well we should see draws from a standard normal distribution); bottom: Simulated ratiometric estimator (with error bars given by the standard error) in black and ’actual’ [Ca^2+^] in red.

The upper part of Fig. 3 is only a qualitative way of checking that the normalized residuals follow a standard normal distribution. A quantitative assessment is provided by the Shapiro-Wilk W statistic, that is here: 0.987; giving a p-value of 0.128. There is therefore no ground for rejecting the null hypothesis that the normalized residuals are IID draws from a standard normal distribution.

As an additional, visual but less powerful test, we plot the empirical cumulative distribution function (ECDF) of the normalized residuals together with the theoretical (normal) one and with Kolmogorov’s confidence bands (Fig. 4). If the empirical ECDF arises from a normally distributed sample with mean 0 and SD 1, it should be *completely* contained in the 95% confidence band 95% of the time and in the 99% band, 99% of the time (these are *confidence bands* not collections of pointwise confidence intervals).

**Figure 4:**
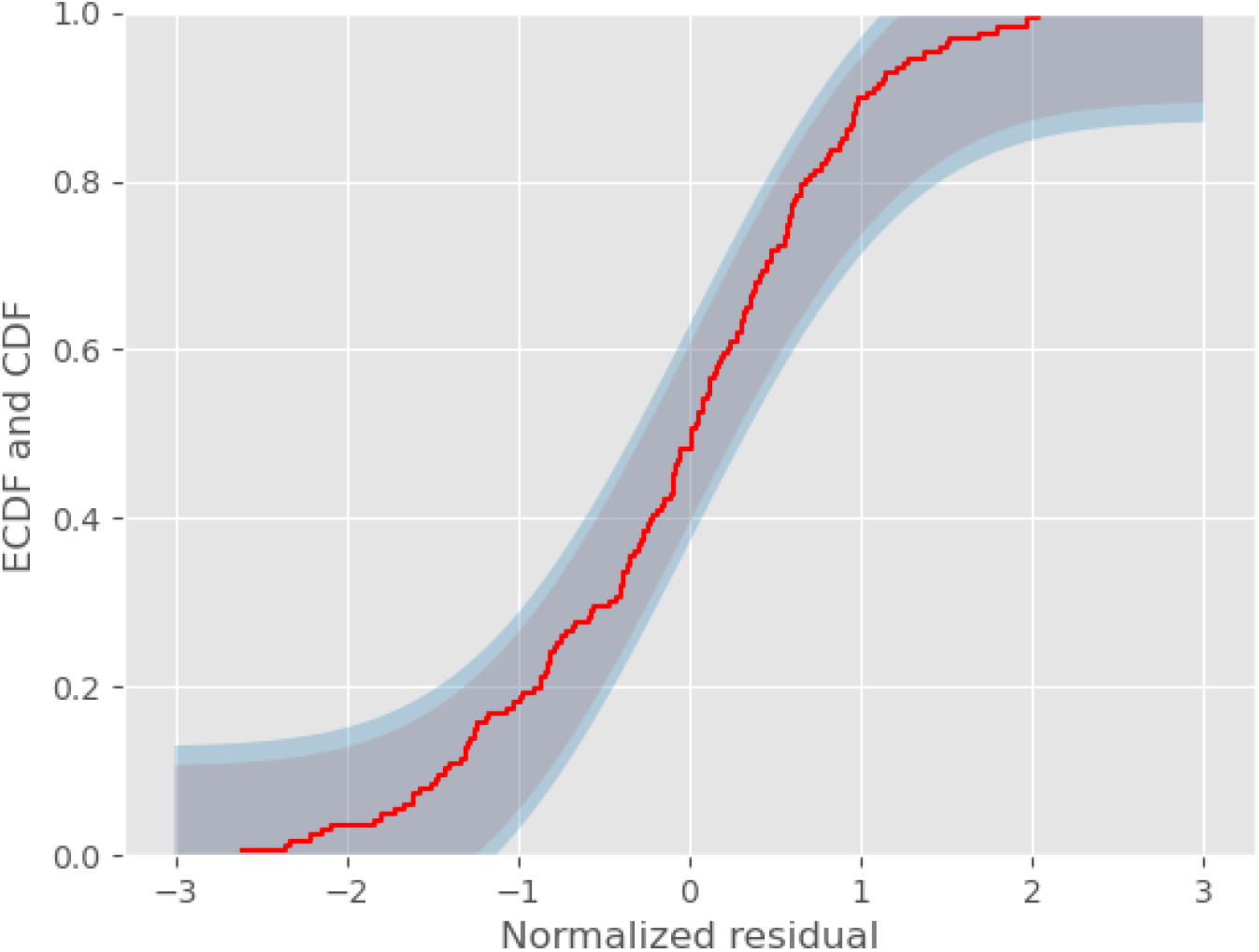
Empirical cumulative distribution function (ECDF) of the normalized residuals (red) together with 95% (grey) and 99% (blue) Kolomogorov confidence bands.

We conclude from these visual representations and formal tests that our normalized residuals follow the expected standard normal distribution, implying that our proposed method for getting the standard errors of the ratiometric estimator is fundamentally correct.

## 6. Discussion

We have presented a new and simple method for getting standard errors on calcium concentration estimates from ratiometric measurements. This method does not require any more data than what experimentalists using ratiometric dyes like Fura-2 are usually collecting: measurements at 340 and 380 nm both within a region of interest and within a background measurement region. Once the errors bars have been obtained, arbitrary models can be fitted to the calcium transients–by weighted nonlinear least-squares [4]–and meaningful confidence intervals for the parameters of these models will follow as illustrated in the companion paper. The present contribution is therefore best viewed a major simplification of the ’direct approach’ of [2]. In contrast to the latter, the new method does not require a ’3 wavelengths protocol’, it does not require either a precise fit of the autofluorescence dynamics at the three wavelengths and is therefore much easier to implement. Such a 3 wavelengths protocol can nevertheless be useful since it allows experimentalists to perform self-consistency checks and to detect potentially wrong calibration parameter (Sec. A). We provide moreover two independent implementations, one in C and one in Python, they are open source and freely available. The rather verbose Python implementation of the heart of the method (Sec. 4.4) requires 25 lines of code and nothing beyond basic numpy functions. We are therefore confident that this method could help experimental physiologists getting much more quantitative results at a very modest extra cost.

## A. A Self-consistency check

## A.1. Plug-in *aduλ* estimates

If we have data where a ’3 wavelengths protocol’ was implemented (that is, flashes at 340, 360 and 380 nm at each time point were made), we can use a self consistency check of the data generation model defined by Eq. 4, 6, 2 and 3. From the 6 measureed sequences:

- {*adu*340(*t*_1_)*, …, adu*340(*t_n_*)},
- {*adu*340*B*(*t*_1_)*, …, adu*340*B*(*t_n_*)},
- {*adu*360(*t*_1_)*, …, adu*360(*t_n_*)},
- {*adu*360*B*(*t*_1_)*, …, adu*360*B*(*t_n_*)},
- {*adu*380(*t*_1_)*, …, adu*380(*t_n_*)},
- {*adu*380*B*(*t*_1_)*, …, adu*380*B*(*t_n_*)},

we compute the sequence of ratiometric calcium estimates 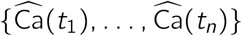. If we also have a way of getting 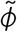 such that:

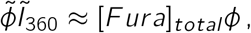

where

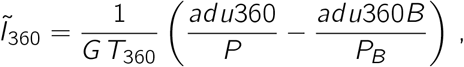

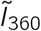 is as estimate of the fluorescence intensity per pixel, per time unit due to Fura (we discuss in the next subsection how to obtain 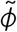), we will get:

- 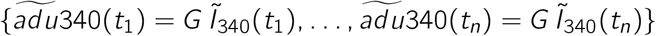,
- 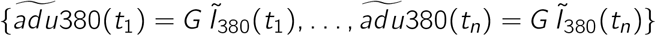

where 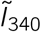 and 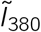 are ’plug-in’ estimates of *I*_340_ and *I*_380_ obtained from Eq. 4 and 6 by plugging in 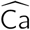:

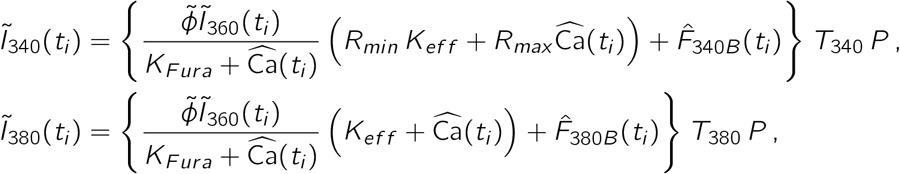

with

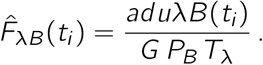

The 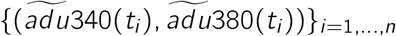 should then be comparable to the original {(*adu*340(*t_i_*), *adu*380(*t_i_*))}_*i*=1*,…,n*_. We have here a self-consistency criterion, not a test since we are using the measured *aduλ(t_i_)*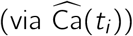 to get 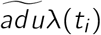.

## A.2. Estimating 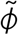

We introduce

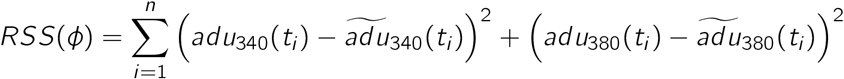

and define 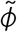 as the argument minimizing *RSS*(*ϕ*). We just need to solve a nonlinear least-square problem.

## A.3. Illustration

As an illustration we compute the plug-in estimators in two situations both based on the first transient of dataset `DA_121219_E1`. In the first case we use the calibrated values for *R_min_* and *R_max_* (left column on Fig. 5). In the second case we use an *R_min_* 20% smaller than the calibrated value and an *R_max_* 20% larger (right column on Fig. 5).

**Figure 5:**
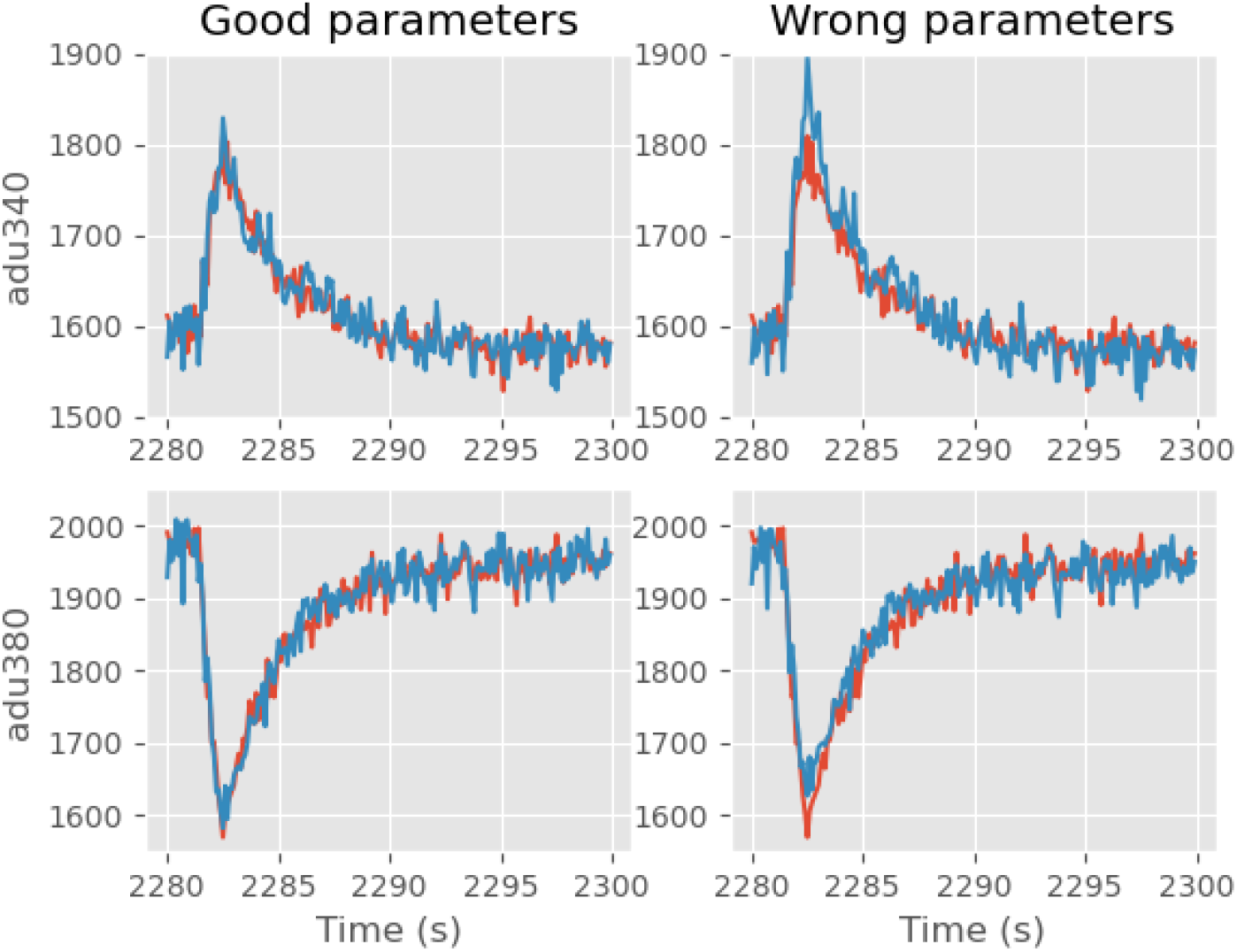
Observed (red) and plug-in (blue) ADU at 340 (top) and 380 nm (bottom) for the first transient. On the left, the calibrated parameters were used while on the right an *R_min_* underestimated by 20% and and *R_max_* overestimated by 20% were used.

We see that the plug-in estimator does indeed provide a self-consistency check (the plug-in estimator values follow closely the observed values on the left side of Fig. 5) and is sensitive to calibrated parameter deviations from their ’true’ values (the peak / valleys ’plug-in’ deviate from the observed values on the right side of Fig. 5).

## B. Propagation of uncertainty

We outline in this section how to reach Eq. 11, 12 and 13 of Sec. 4.4.1. We first need to remember that is *X* and *Y* are two *independent* random variables with mean 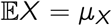 and 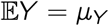 and variance 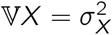 and 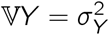, then if *Z* = *a* + *bX* + *cY* (*a, b, c* ∈ ℝ) we have:

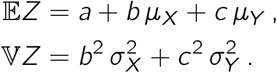

Eq. 11 is a direct consequence of the last equality. If *X* is (approximately) normally distributed with 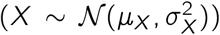 as well as *Y*, we can write: *X* ≈ *µ_X_* + *Z*_1_*σ_X_* and *Y* ≈ *µ_Y_* +*Z*_2_*σ_Y_*, where *Z*_1_ and *Z*_2_ are independent and follow a standard normal distribution, 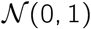. If now *Z* = *f* (*X, Y*) and the partial derivatives of *f* at (*µ_X_, µ_Y_*) exist then:

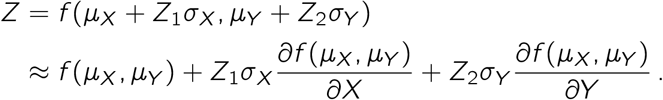

This is just a first order Taylor expansion and that is where the ’small enough standard error’ assumption is necessary. *Z* is then (approximately) a linear combination of two independent standard normal random variables and we immediately get:

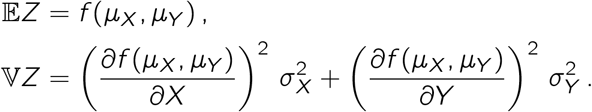

Eq. 12 follows directly by computing the necessary partial derivatives, while Eq. 13 requires the computation of a single derivative.

## C. Auto-fluorescence dynamics

## C.1. General features

The evolution of the *aduλB* is shown on Fig. 6. We see that the autofluorescence runs down when high frequency flashes are applied during the 3 transients, with a partial recovery between transients.

**Figure 6:**
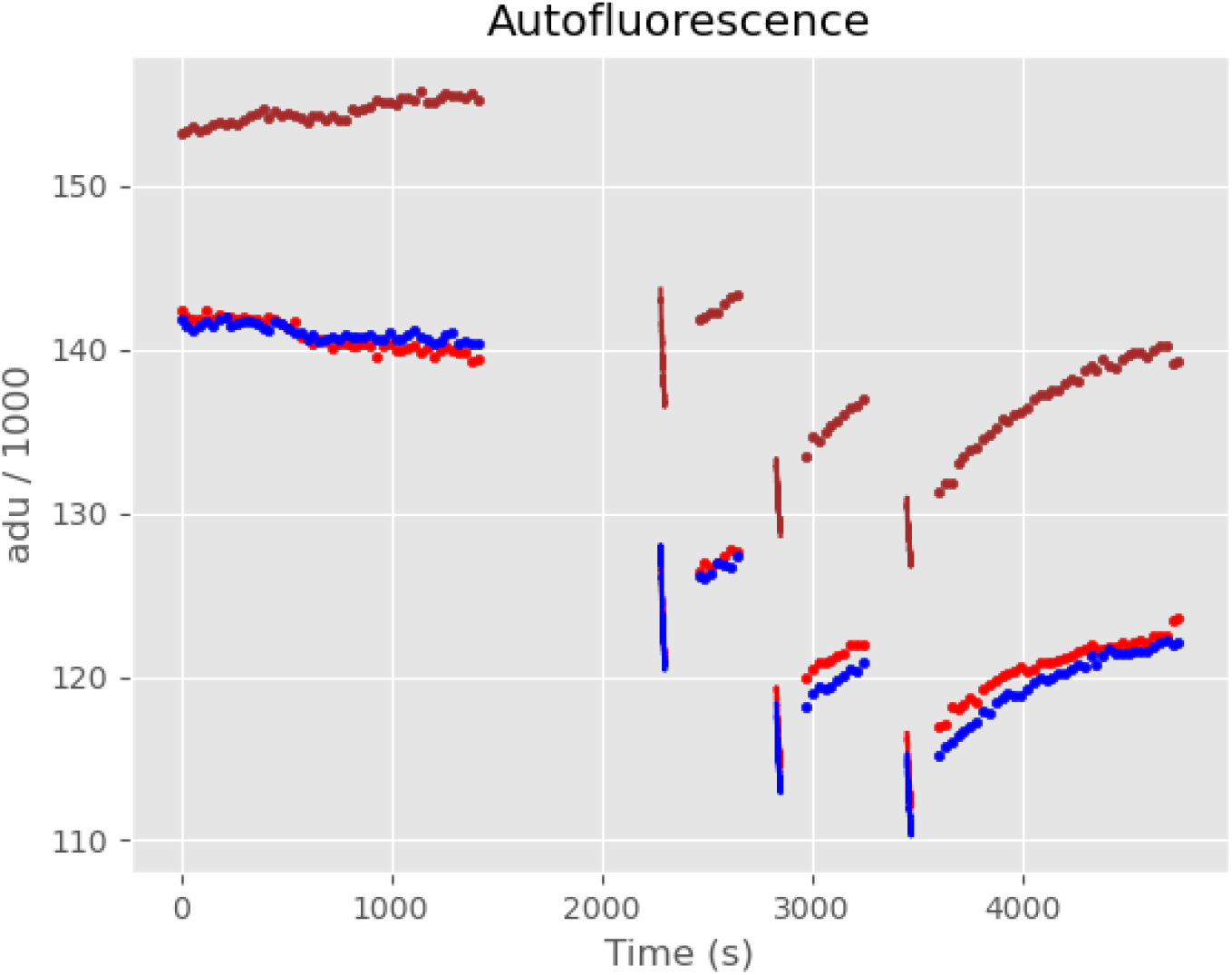
Autofluorescence at 3 excitation wavelengths, 340 nm (red), 360 (blue), 380 (brown). Both during low frequency stimaltions (four portions made of dots) and during the transients where a higher frequency stimulation was applied (3 groups with almost vertical lines).

## C.2. Within transient dynamics

The ’direct method’ of [2] requires the knowledge of the autofluorescence value at each time point during a transient at both 340, 360 and 380 nm, since Eq. 4 and 6 are fitted directly to the recorded adu340 and adu380 and they depend on the total Fura concentration at transient time that is estimated from the difference of the 360 nm measurements in the ROI and the BMR. We therefore take a closer look a the autfluorescence dyanmics during the first transient (Fig. 7).

**Figure 7:**
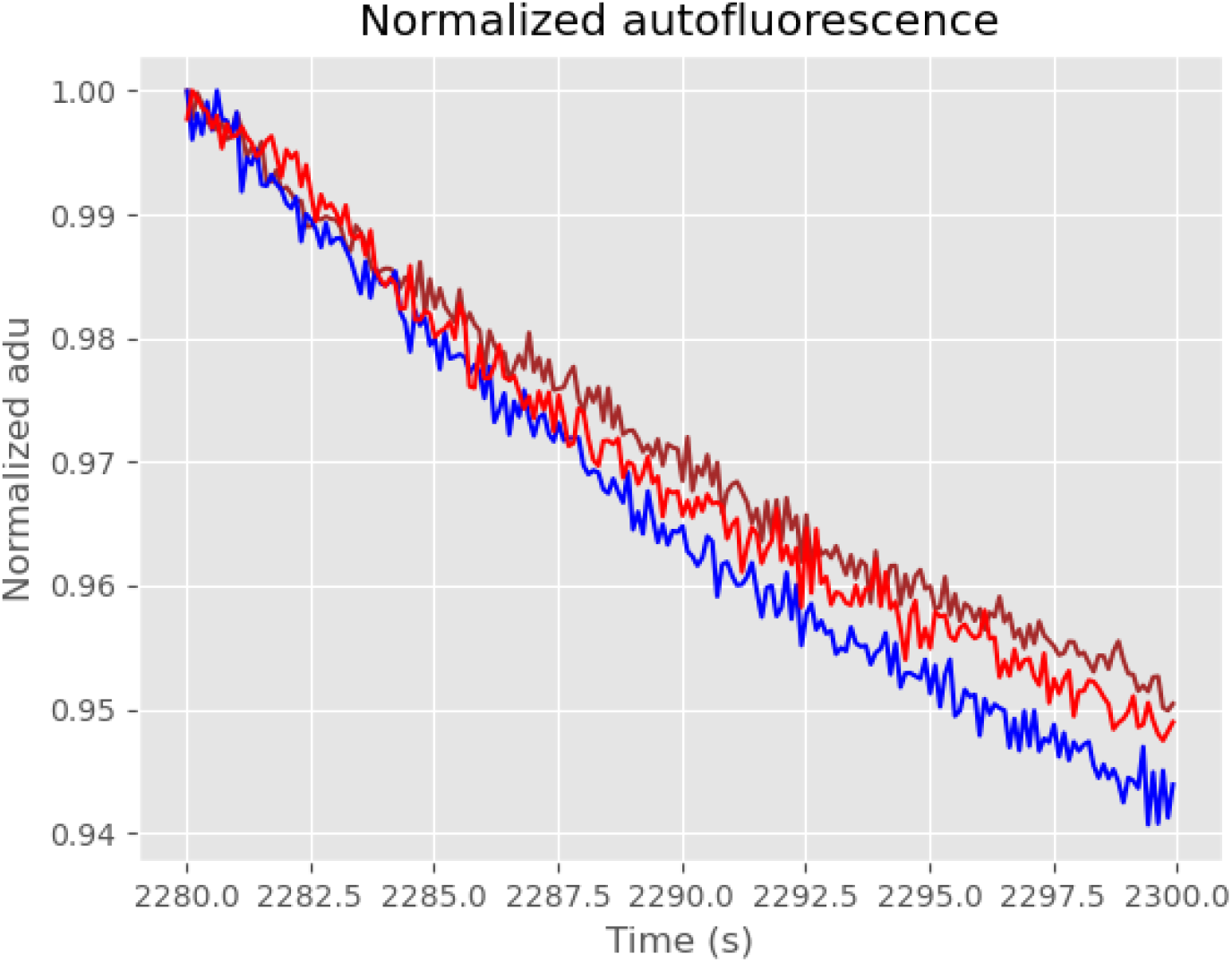
Normalized autofluorescence at 3 excitation wavelengths, 340 nm (red), 360 nm (blue) and 380 (brown) during the first transient. At each wavelength, the normalization is performed by dividing each value by the maximal one.

At that stage we can fit a straight line plus a cosine function whose period is the duration of a transient. That’s a good way to capture the main structure in the transient, but is still does not account for the full signal variability (Fig. 8). As can be seen from the normalized residuals–the residuals divided by the standard deviation–that should be very nearly independent random draws from a standard normal distribution if the model is correct, there are finer structures left (like the double valley on the 380 nm residuals) meaning that those fits won’t pass formal goodness of fit tests. Indeed if we apply Pearson’s *χ*^2^ tests to these stabilized residuals we get:

**Figure 8:**
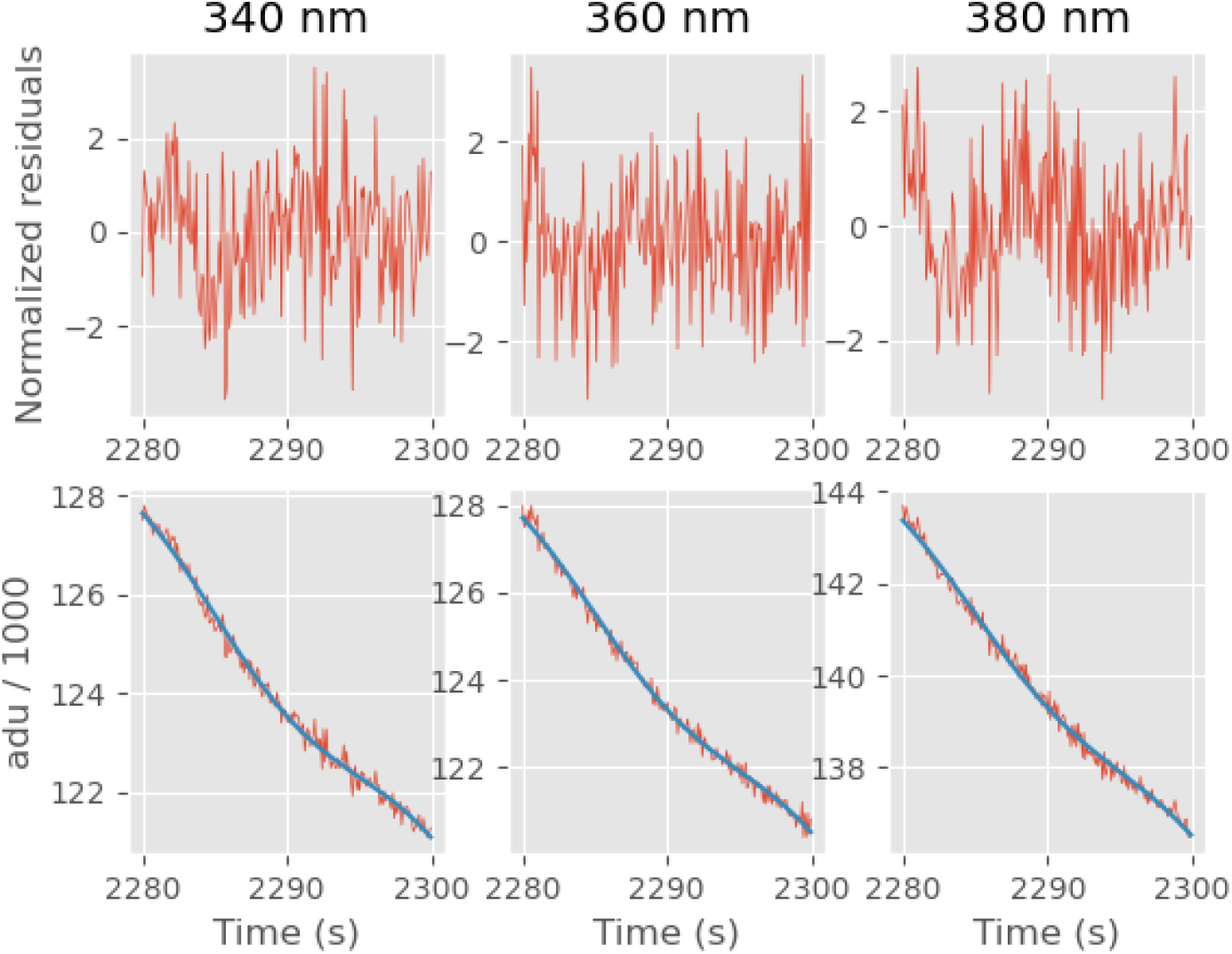
Bottom: Autofluorescence (red) at 340 nm (left), 360 nm (middle) and 380 nm (right) together with a straight line plus cosine function fit (blue). Top, the normalized residuals: 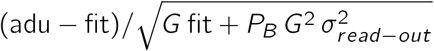.

- at 340 nm a residual sum of squares (RSS) 326, leading to a 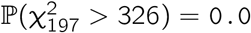,
- at 360 nm a RSS of 288, leading to a 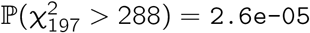,
- at 380 nm a RSS of 275, leading to a 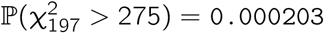,

We are then left with three possibilities:

1. try to refine the ’straight line plus cosine function’ empirical model in order to get acceptable fits,
2. try to get a better understanding of the autofluorescence dynamics,
3. find another way to get error bars on our estimates.

Since we wanted to propose an ’as general and easy as possible’ method we chose the third approach in the present manuscript.

## D. Summary figure

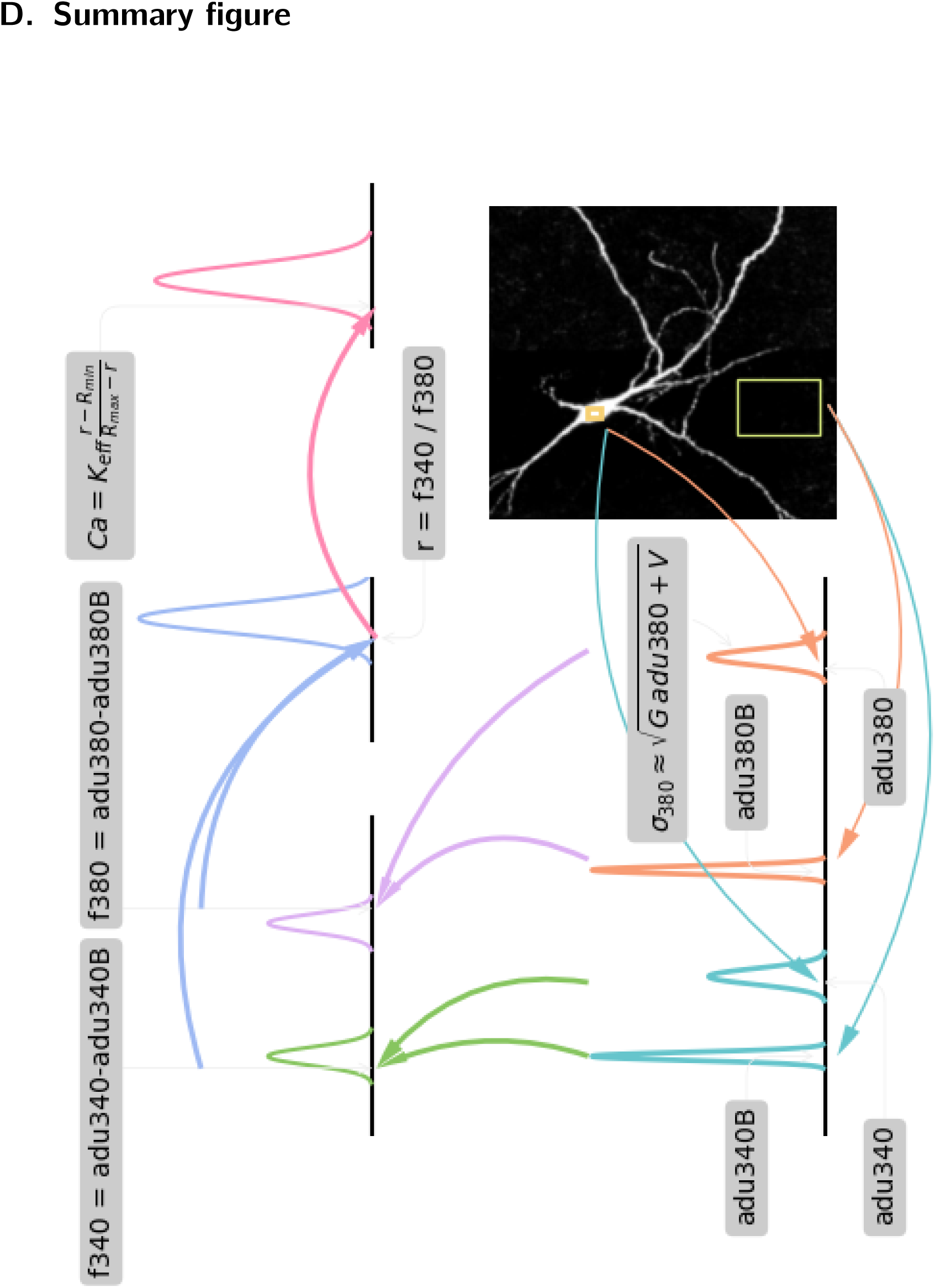

